# Detecting Differential Alternative Splicing in Mass Spectrometry-based Proteomics Data

**DOI:** 10.1101/2023.09.19.558203

**Authors:** Constantin Ammar, Gergely Csaba, Armin Hadziahmetovic, Catherine G. Vasilopoulou, Markus Gruber, Matthias Mann, Ralf Zimmer

## Abstract

Alternative splicing can substantially diversify biological cell states and influence cellular function. The functional impact of splicing has to be estimated at protein level, typically by mass spectrometry (MS) -based proteomics. Although this technology measures increasingly large peptides sets, distinguishing isoform-specific peptides are rare, limiting detection and quantification of splicing. We introduce MS-EmpiReS, a quantification-based computational approach for differential alternative splicing detection in proteomics data. Its core principle is to differentially quantify peptides mapping to different regions of genes. This approach increased the number of testable peptides hundred-fold in a clinical cancer cohort, resulting in a large number of cancer-relevant splicing candidates. Splicing events detected by both MS-EmpiReS and deep RNA sequencing correlated well but also provided complementary information. The proteomics data allowed us to define a per-sample splicing score to separate cancer conditions. Finally, deep brain proteomes from different mice separated strongly by the lower abundance protein splicing isoform.

## Introduction

The exon-based gene structure of eukaryotic organisms enables the production of multiple proteins from a single gene via alternative splicing (AS). The impact of AS on the proteome is controversial, as it often remains unclear whether and to what extent the alternative transcript observed in deep sequencing data will actually result in relevant amounts of alternative protein products (*isoforms*) ^1–6^. Alternative splicing on the protein level is usually detected in mass spectrometry (MS) data by utilizing the sequence information of identified peptides. For example, junction peptides can be identified which could span over a spliced-out exon. If an additional peptide within the exon is identified, both together are a clear indication of alternative splicing. Intron retentions and alternative start sites can also be identified via the respective peptides. Such *sequence-exclusive* approaches^3,5,7,8^ are the most basic form of AS detection, usually aiming at the assessment of the general prevalence of splicing, for example in the human proteome.

In order to obtain detailed insights into the regulation of splicing, it is necessary to not only assess whether alternative isoforms are present or not, but also to quantify differences in the expression of these isoforms. This quantitative aspect has been less studied in proteomics, as usually only a very small fraction of isoform-specific peptides are sequence-exclusive^5,9–11^. To address this, one could test whether one isoform of a gene changes differently as compared to another isoform of the same gene, a process termed *differential alternative splicing* (DAS). While several tools exist that detect quantitative variations of peptides^12–19^, there is currently no tool for the systematic screening and statistical evaluation of proteomic DAS, analogous to the tools available for transcriptomics data^20–24^. In the following study, we introduce a novel computational method to investigate quantitative proteomic DAS over differing biological conditions. It is based on our differential quantification method for “Mass Spectrometry analysis using Empirical and Replicate bases statistics” (MS-EmpiRe)^25^. MS-EmpiRe relies on peptide fold-changes that enable precise fold-change estimations (0.1-2 fold errors)^25,26^. It then utilizes empirical between-replicate distributions to assign probabilities to individual peptide fold-changes. In our new extended algorithm we present a framework to compare these peptide fold-changes against each other in the context of splicing. The ‘MS-EmpiReS’ algorithm enables us to score whether peptides mapping to one region of the protein have significantly different fold-changes from peptides mapping to another region. In combination with isoform mappings from the Ensembl^27^ database, we assemble these regional fold-changes to fold-change differences between isoforms and thereby identify candidates for all types of DAS. We go on to show that this quantitative approach enables order-of-magnitude improvements in the number of testable peptides as compared to the sequence-exclusive approach.

After benchmarking MS-EmpiReS on simulated data and on *E. coli*, which does not have splicing, we apply it to a cohort of one hundred colon cancer patients. Testing for DAS in about 3000 genes detected 150 DAS events in the data, much more than the sequence-exclusive approach. Compared to RNA-seq, we find both a stable subset of isoform ratios as well as complementary information, and the proteomics data can be used to generate a splicing score which discriminates cancer from non-cancer tissue. In deep proteomes of mouse brains, we detect individual-specific splicing patterns driven by the lower abundant isoform.

MS-EmpiReS is available under https://www.bio.ifi.lmu.de/software/msempire_s/index.html. The tool performs DAS analyses on standard peptide input tables and provides differential splicing tables that can be accessed through a web browser. It also provides detailed on-demand visualisation of individual splicing results using simple commands and can be run on a standard laptop with processing times in the order of tens of minutes.

## Results

### A model for DAS detection based on parameter-free empirical distributions

Figure 1A visualizes the possible cases of splicing regulation between two biological conditions. These are: (i) switching of isoforms between conditions (top), (ii) expression of an additional isoform in one condition (middle) and (iii) differential abundance changes between two expressed isoforms in both conditions (bottom). It is challenging to distinguish between non expressed and non detected peptides in MS proteomics data^7^, which would be necessary to identify case (i), therefore we focus on cases (ii) and (iii) in the following.

**Figure 1.**
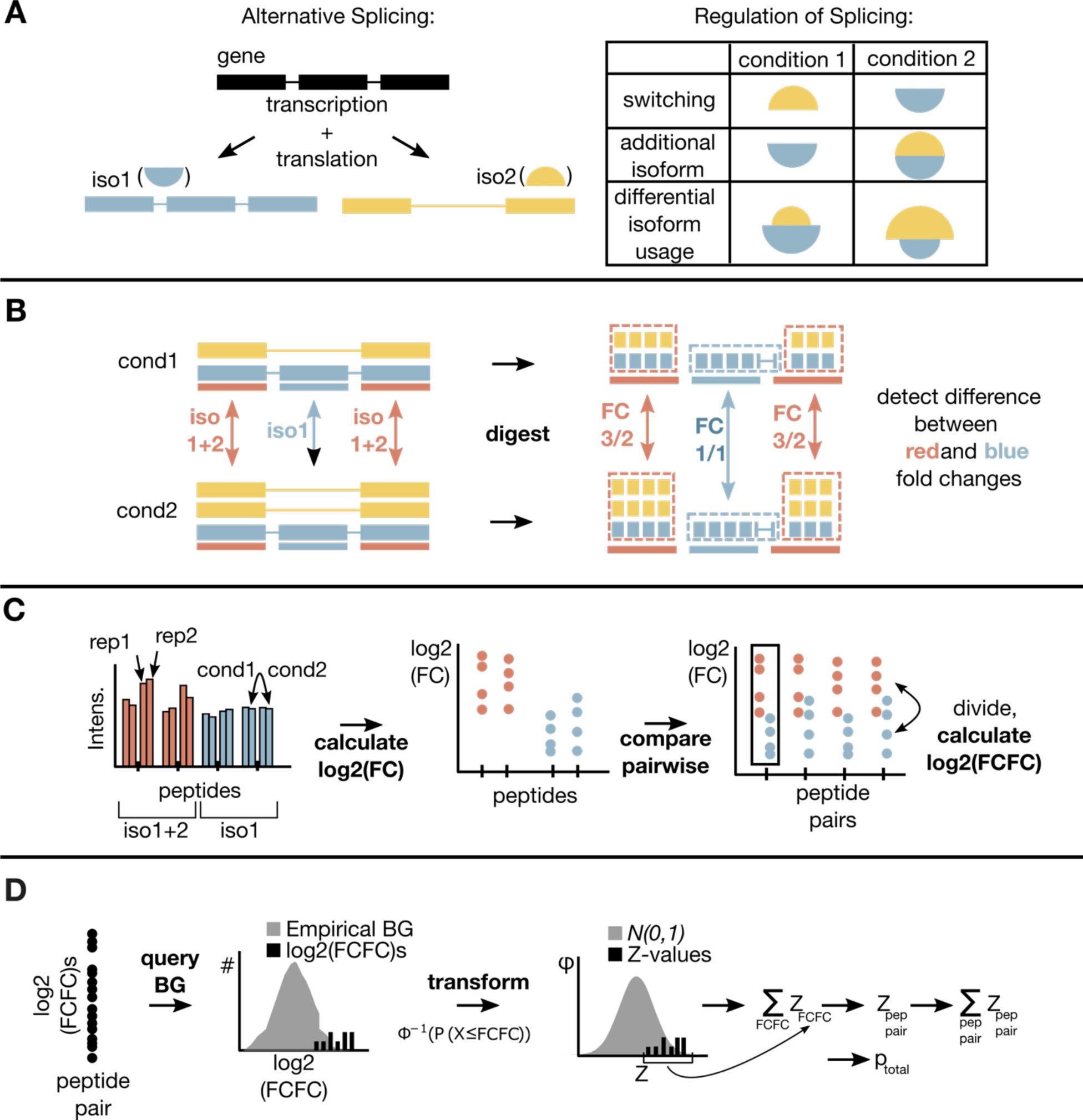
MS-EmpiReS workflow. A) Exemplary alternative splicing event (exon skipping) and and different forms of splicing regulation. B) Principle of quantitative splicing detection for differential isoform usage. The yellow isoform doubles in condition 2. Peptides resulting from enzymatic digestion (small squares) either map only to iso1 (marked blue) or both iso1+2 (marked red). The fold-changes of the red peptides 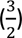 are different to the fold-changes of the blue peptides 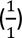, which can be detected by MS-based proteomics. The yellow isoform has no exclusive region and thus no unique peptides in this example. C) Peptide level comparison between red and blue regions, with an example of two red and two blue peptides with two replicate samples each. Peptide fold-changes between conditions are assessed in a first step. Red and blue peptides are then compared in a pairwise manner and “fold-changes of fold-changes” (FCFCs) are assessed. D) The log2(FCFC)s are used to query an empirical error distribution which is derived from replicate measurements. The observed log2(FCFC)s can be expressed as Z-values (with direction of the change), summarized and then transformed to a p-value.

We illustrate the basic principle of MS-EmpiReS with an example of two expressed isoforms in two conditions (Figure 1B). Isoform 1 doubles from condition 1 to condition 2, while isoform 2 does not change. The peptides resulting from enzymatic digestion either map only to isoform 2 (blue) or to both isoform 1 and isoform 2 (red). Note that peptides mapping only to isoform 1 or peptides mapping to an additional isoform may also exist. These scenarios can always be reduced to a similar case as displayed here and are hence omitted in the figure for clarity. The fold-changes of the red peptides should center around 3/2, because there are two copies in condition 1 and three copies in condition 2. The fold-changes of the blue peptides should center around 1, because there is no change in isoform 2. With MS-EmpiReS, we statistically evaluate the fold-change differences between such groups of peptides as described below. We additionally complement this quantification-based approach with a sequence-exclusive approach to utilize the full information available in the dataset.

The bioinformatics pipeline of MS-EmpiReS (see Methods) is depicted in more detail in Figure 1C with the simple example case of two peptides in each group and two replicate measurements for each peptide. The fold-changes between conditions are determined for every peptide and all peptide pairs between the two groups are formed. For every peptide pair, four against four peptide fold-changes are compared and the fold-changes are divided, resulting in 16 “fold-changes of fold-changes” (FCFCs) blue/red, which are log2 transformed. The absolute value of the log2(FCFC) indicates how dissimilar the change between isoforms is. Positive log2(FCFC)s reflect that the blue group changes more strongly than the red group. The log2(FCFC)s are compared to an empirical error distribution, describing the log2(FCFC)s of non-changing peptide pairs. From the empirical error distribution and a given log2(FCFC), a normally distributed Z-value can be derived. The Z-values are combined using a modified Stouffer^28^ approach to calculate an overall score, which tests the null hypothesis: no difference in the change of the two peptide groups^25^. We denote the multiple testing corrected score as *p_adj_*. Dependencies of the variables have to be taken into account at several points of the calculation. Peptides are mapped to protein isoforms based on the Ensembl genome annotation^27^. To detect quantitative differences between isoforms we use the log2(FCFC)s of peptide pairs, where the peptides stem from different Ensembl annotated isoforms. These contain the current state of the art of known Isoforms comprising all relevant splice events (exon skippings, alternative donor/acceptor sites, intron retentions, etc.)^29,30^ .

As some genes have a large number of annotated isoforms not all of which are expressed as proteins in the condition under study, we determine *equivalence classes* that group peptides unique to a specific (set of) isoforms (see Methods). For simplicity we will refer to the equivalence classes as isoforms. Thereby, log2(FCFC)s are compared for peptide pairs relevant for the isoforms of interest. Again, log2(FCFC)s are accumulated over all these relevant peptide pairs in order to estimate the significance of DAS.

### Benchmarking

#### Randomly sampled intensities result in uniformly distributed p-values

For a first benchmark, we tested if our noise model fulfils the basic requirements of statistical models. We simulated proteins with randomly sampled intensities, representing non-spliced proteins (see Methods). For each protein, we randomly distributed the corresponding peptides into two groups, representing two virtual isoforms (i.e. “red” and “blue” peptides in Figure 2A, left) and tested them against each other with MS-EmpiReS. Each group had to contain at least two peptides. As there is no systematic shift in the data, the p-values from the tests should be uniformly distributed. The data confirmed this to be the case, also when one isoform has few peptides or replicates and the other isoform has many peptides or replicates (Figure 2A, right).

**Figure 2.**
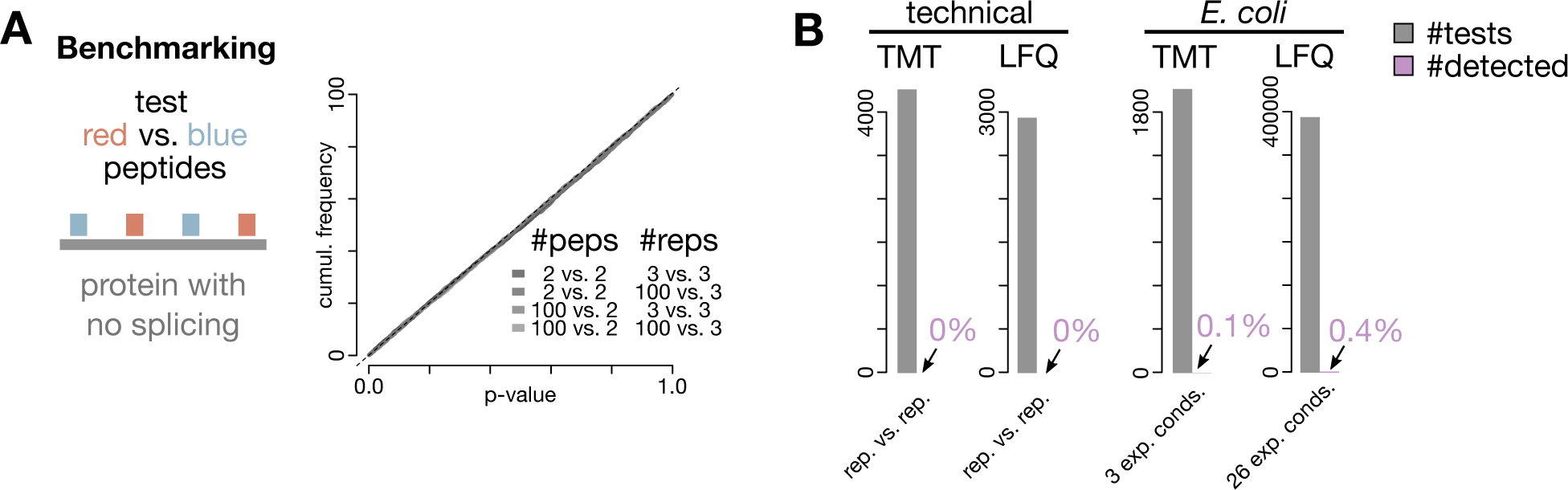
Benchmarking. A) Simulated peptides of non-spliced proteins were randomly assigned to two groups and tested against each other. Simulating peptides by drawing random intensities resulted in uniform p-values as required, even for drastically different peptide and replicate numbers. B) Negative controls on several experimental (LFQ and TMT) datasets using technical replicates and the non-splicing organism E. coli (such that every hit is a false positive). We applied MS-EmpiReS to random subsets of peptides and displayed significant hits (adjusted p-value (p_adj_) < 0.01) in violet.

#### Testing on experimental datasets with no splicing shows well controlled false positives

As a next step we benchmarked MS-EmpiReS for false positives in situations where there is no regulated splicing, and where a positive hit would indicate a false positive. First, we tested MS-EmpiReS on technical replicate datasets for both data dependent acquisition label free quantification (LFQ) and Tandem Mass Tag (TMT) data (ref1, ref2). To simulate differential conditions, we split six replicate measurements of human cell lysate into two groups and tested them against each other. Testing thousands of proteins, MS-EmpiReS did not report single significant hit (< 1% *p_adj_*) (Figure 2B, left). This indicates excellent handling of variation and biases within a proteomics replicate workflow.

Finally, we compared quantitative *E. coli* proteomics datasets (three conditions for TMT and 27 conditions for LFQ data) as prokaryotes have no splicing mechanism. This dataset was particularly challenging because of extensive proteome remodelling between conditions (differential regulation for ≈30% of the proteome on average), thereby introducing both technical and biological differences that could be mistaken for DAS. After testing, we indeed observed 0.4% false positive hits. Inspection showed that some peptides had systematic shifts that were consistent between replicates (Supplemental Figure 1). This might be due to slight differences in sample preparation per condition. As our model explicitly tests for shifts between groups of peptides, combinations of such peptide shifts could lead to significant hits. However, as the rate of such events was low at 0.4% we deemed our model sufficient for confident splicing identification.

### Analysis of a clinical proteomics dataset

#### MS-EmpiReS enables comprehensive screening of thousands of genes for splicing

Having established basic performance characteristics of MS-EmpiReS, we next asked if it could detect cancer associated DAS. We selected a clinical proteomics cohort of about 100 colon cancer patients, measured by the Clinical Proteome Tumour Analysis Consortium (CPTAC)^31^. In that study, two samples had been extracted from each patient, one from cancerous tissue and one from normal-appearing adjacent tissue (termed ‘normal’ in the following). We first detected sequence-exclusive splicing with MS-EmpiReS (see Methods).

We then filtered out peptides with less than five measured patients in cancer or normal, resulting in about 138,000 testable peptides, which we quantitatively tested with MS-EmpiReS (Figure 3A, left). Each protein with at least two precursors in at least two equivalence classes is amenable to DAS testing, which resulted in around 3,200 testable genes in this CPTAC dataset, 50 times more than in the sequence-exclusive approach (Figure 3A, middle). This first quantitative proteome-wide screening for DAS resulted in six-fold more cancer associated splicing events than with the previous sequence-exclusive approach.

**Figure 3.**
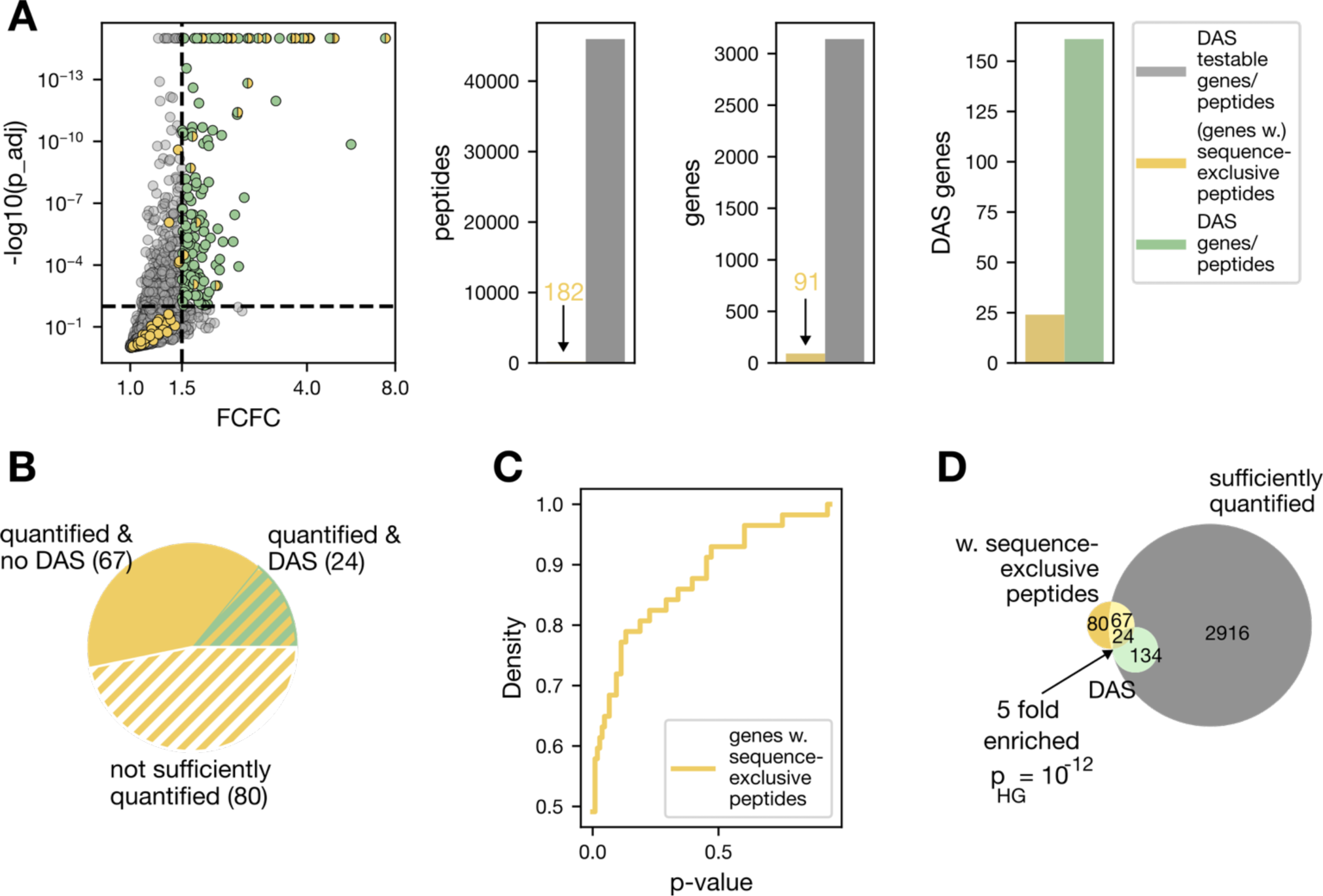
Application of MS-EmpiReS to a CPTAC colon cancer data set of about hundred patients^31^. A) The results of isoform changes are displayed as a volcano plot with absolute FCFCs. Genes containing isoform pairs with *p_adj_*<1% and FCFC>1.5 are classified as DAS (green). The yellow color indicates genes with sequence-exclusive peptides, where both isoforms were sufficiently quantified. A large number of these is in the insignificant area of the volcano plot. Counts underlying the volcano plot are displayed on the right. The number of testable genes is increased by one and that of peptides (grey) by two orders of magnitude compared to the sequence-exclusive approach resulting in a six-fold increase of the number of DAS genes. B) Overview over genes with sequence-exclusive peptides. A majority of these was not accessible to DAS assessment because they had too few data points. C) Cumulative distribution of p-values of genes with sequence-exclusive peptides. Around 50% of these genes have insignificant p-values >0.01 (before multiple testing correction). D) Genes with sequence-exclusive peptides are strongly enriched in the DAS genes quantitatively classified by MS-EmpiReS.

#### The sequence-exclusive approach fails to comprehensively quantify DAS

We detected 171 genes to be spliced based on the sequence-exclusive approach. Less than half of these genes with sequence-exclusive peptides are properly quantified (Figure 3C), either because there are insufficient measurements or not enough measured peptide precursors to ensure proper quantification (see Supplemental Table 1). Of those that were properly quantified, 50% had no significant p-value for regulation in cancer even before multiple testing correction (Figure 3B, Supplemental Figure 2). Therefore, mere detection of isoform-specific peptides without additional quantitative assessment is not sufficient to obtain reliable information about splicing regulation. For more consistent quantification, targeted data acquisition methods might be necessary^32,33^ in the sequence-exclusive approach.

#### The quantification-based approach is consistent with the sequence-exclusive approach

Although the sequence-exclusive approach alone does not provide comprehensive information, it provides useful additional information to validate and complement our quantitative approach. In particular, genes that are detected as DAS using the quantitative approach should be more likely to contain sequence-exclusive peptides as compared to randomly selected genes. Indeed, we found a five-fold enrichment (p < 10^-12^, Figure 3D), of genes with sequence-exclusive peptides within the DAS genes, indicating that our quantification-based approach is suitable to detect actual splice events. Next, we checked 20 of the top ranked DAS genes (Table 1) against a recent database on splicing in the human proteome, which was generated with a sequence-exclusive approach using large scale profiling of MS proteomics data of human tissues^5^. This revealed that 75% of them had prior MS-based evidence of splicing. Furthermore, we found literature-based evidence for splicing in the cancer context for the remaining five genes and altogether for 19 of the 20 top ranked DAS genes (Supplementary Table 3).

**Table 1.** Twenty of the top ranked DAS genes in the CPTAC data set (see Supplemental Table 2 for full list). The “Splicegene Lau et al.” column indicates whether the gene is listed as alternatively spliced in the protein splicing database^5^. The “Literature Cancer Splicing” column indicates whether there are explicit literature mentions of the gene as being alternatively spliced in the context of cancer (Supplemental Table 3). Bold genes were shown in detail in Figure 3.

| Gene | Protein Name | SpliceGene Lau et al. | Literat. Cancer Splicing |
| --- | --- | --- | --- |
| ACTN1 | actinin alpha 1 | y | y |
| ACTN4 | actinin alpha 4 | y | y |
| CALD1 | caldesmon 1 | y | y |
| CAPZB | capping actin protein of muscle Z-line subunit beta | n | y |
| CFL1 | cofilin 1 | n | y |
| CHID1 | chitinase domain containing 1 | y | n |
| COL6A3 | collagen type VI alpha 3 chain | y | y |
| EPB41L2 | erythrocyte membrane protein band 4.1 like 2 | y | y |
| H2AFY | macroH2A.1 histone | y | y |
| LRRFIP1 | LRR binding FLII interacting protein 1 | y | y |
| **MAP3K20** | mitogen-activated protein kinase kinase kinase 20 | n | y |
| PDLIM5 | PDZ and LIM domain 5 | y | y |
| PDLIM7 | PDZ and LIM domain 7 | y | y |
| **PKM** | pyruvate kinase M1/2 | y | y |
| RPS7 | ribosomal protein S7 | n | y |
| SPTAN1 | spectrin alpha, non-erythrocytic 1 | y | y |
| TNC | tenascin C | y | y |
| **TPM1** | tropomyosin 1 | y | y |
| TPM2 | tropomyosin 2 | y | y |
| TPM4 | tropomyosin 4 | n | y |

#### Examples of clinically relevant alternative splicing candidates

Detailed visualizations of selected DAS events are given in Figure 4 and for all in Supplemental File 1. Tropomyosin 1, a gene that stabilizes smooth muscle actin and regulates its function, has been reported to be a tumour suppressor gene with splice events impacting colony formation and regulatory activity^34^. We see downregulation of both isoforms in cancer, about ten-fold for the *CRA*_*f* isoform and 2-fold for the *CRA*_*c* isoform. Even though the regulation goes into the same direction, MS-EmpiReS clearly resolves the splice event (*p_adj_ <* 10^−15^), potentially indicating a higher functional relevance of *CRA*_*f.* Another example is MAP3K20, also known as ZAK kinase, a member of the MAPKKK family of signal transduction molecules that activates cancer-related signaling pathways such as NF-*κ*B, Wnt/*β*-catenin, and AP1. The ZAK long form (ZAK-LF) and the ZAK short form (ZAK-SF) differ strongly with the ZAK-LF inducing tumour growth in immunodeficient mice^35^. Accordingly, we see a switching event in colon cancer, with the tumour associated isoform being upregulated and the ZAK-SF being downregulated. This indicates a cancer relevant splicing induced signalling switch at the protein level. As we have peptides mapping to isoform 1, isoform 2 and shared peptides between both isoforms, we can estimate the stoichiometries of the isoforms (Supplemental Text T1). This estimation indicates that the ZAK-SF is almost two orders of magnitude more abundant than the cancer up-regulated ZAK-LF, providing crucial information for interpreting functional impact. As a third example the pyruvate kinase M 1/2 gene, which mediates the last step of glycolysis, namely the dephosphorylation of phosphoenolpyruvate to pyruvate is an essential metabolic gene that has been widely studied in the context of cancer^36^. For example, it has been reported that switching of the PKM2 isoform to PKM1 reverses the Warburg effect in cancer cells^37^. Interestingly, we see a slight upregulation of the PKM2 associated peptides but also a stronger downregulation of the PKM1 associated peptides in the patient data.

**Figure 4.**
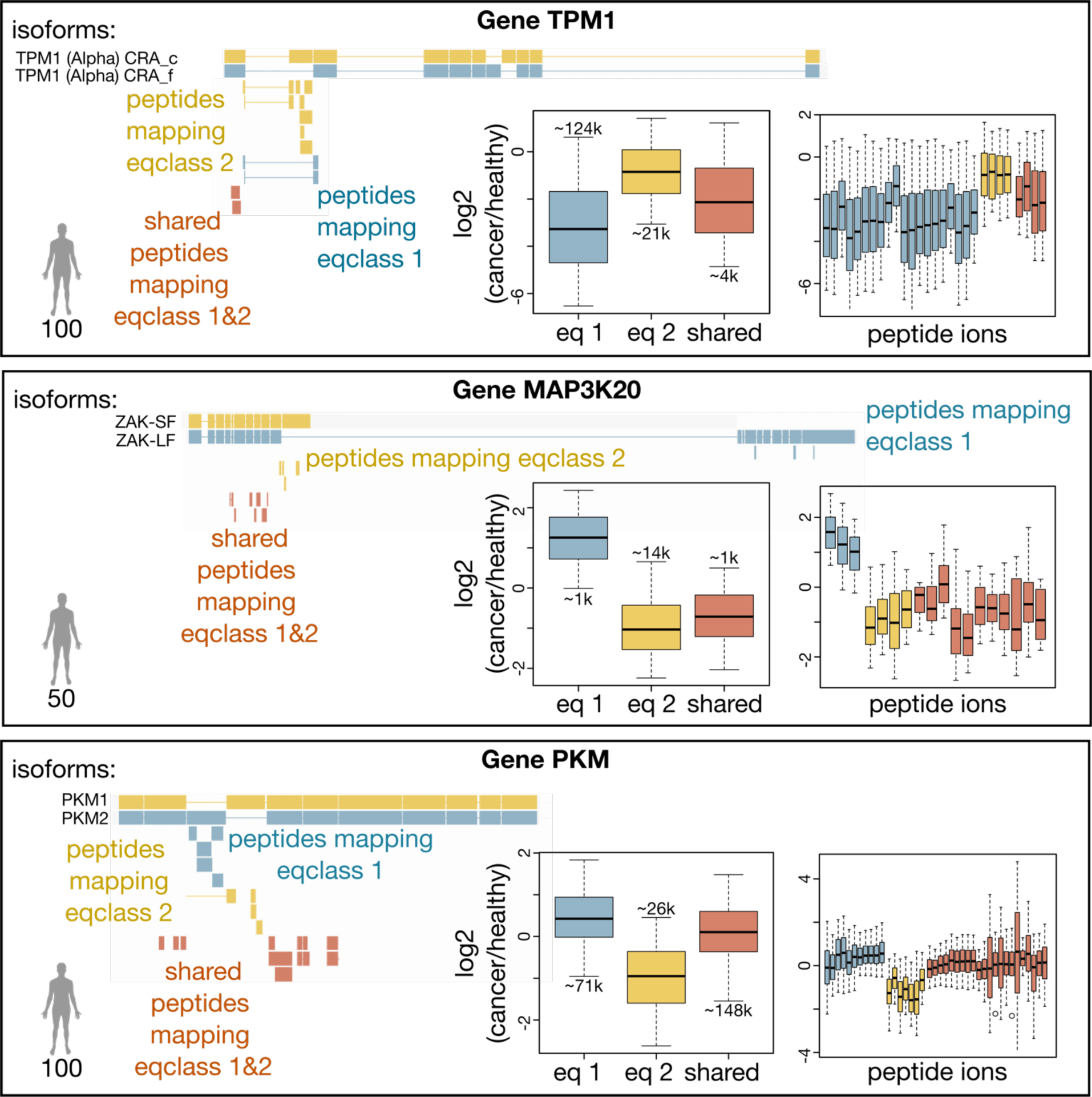
Visualizations of DAS events for TMP1, MAP3K20 and PKM, top scoring genes with important regulatory functions. Two isoforms each are displayed in yellow and blue with quantified peptides aligned below (as in Figure 1). The box plots show fold-changes between normal and cancer for peptides mapping exclusively to the isoforms (blue and yellow), and shared peptides (red). Numbers atop the boxes represent the total underlying datapoints. The numbers in the bottom left corners indicate the quantified patient samples for the DAS event. Plots created by MS-EmpiReS and visually re-arranged. Boxplot elements correspond to the following quantiles: 0.05, 0.25, 0.5, 0.75, 0.95.

### Comparison to RNA-seq data

#### DAS genes for RNA-seq and proteomics diverge even for high-confidence hits

With a detailed and in-depth proteomic cancer associated splicing event in hand, we next compared our data with transcriptomics (RNA-seq) data from an independent clinical cohort (20 patients) from the The Cancer Genome Atlas (TCGA)^38–40^. To this end, we employed our previously described EmpiReS package for the detection of DAS on the transcriptome level^41^ (see Methods). This resulted in around 9000 tested genes, containing more than 90% of those that we had also tested in the CPTAC proteomics data. For each gene, we used the two most significant isoforms in the transcriptomics data as the reference pair, to which we mapped the peptides from the proteomics data (see Methods). This resulted in around 1,800 genes that could be compared (Figure 5A). In this intersection there were 52 DAS events at the transcript level and 97 at the proteomics level (*p_adj_ <* 0.01 and log2(FCFC)>0.5), with an overlap of only 11 genes (Figure 5B). When further restricting to proteins that also had sequence-exclusive peptides, only 5 of 13 genes were classified as DAS in the transcriptomics data (Figure 5C). This divergence indicates either false negatives in the transcriptomics measurements or differences in translation of the isoforms. To further investigate the apparent divergence of DAS in the transcriptome and proteome, we analyzed all genes with DAS on either level. The fold-changes of the individual isoforms from normal to cancer as transcript and as protein have Pearson Correlation Coefficients of around 0.7 to each other (Figure 5D). Given that the transcriptome data is from a different cohort and measurement than the proteome data, this appears to be a relatively high agreement. However, the correlation of fold changes of the isoforms of each gene – the log2(FCFC) – is much lower (0.3), consistent with the fact that the log2(FCFC)s are a more fine-grained measure and dependent on good quantification of both isoforms. This value improves up to 0.76 when only considering DAS genes of proteomics or transcriptomics (Figure 5E, Supplemental Figure 3). Thus, an increase in measurement quality leads to higher log2(FCFC) correlation, indicating that much of the divergence seen is due to limitations of the measurements.

**Figure 5.**
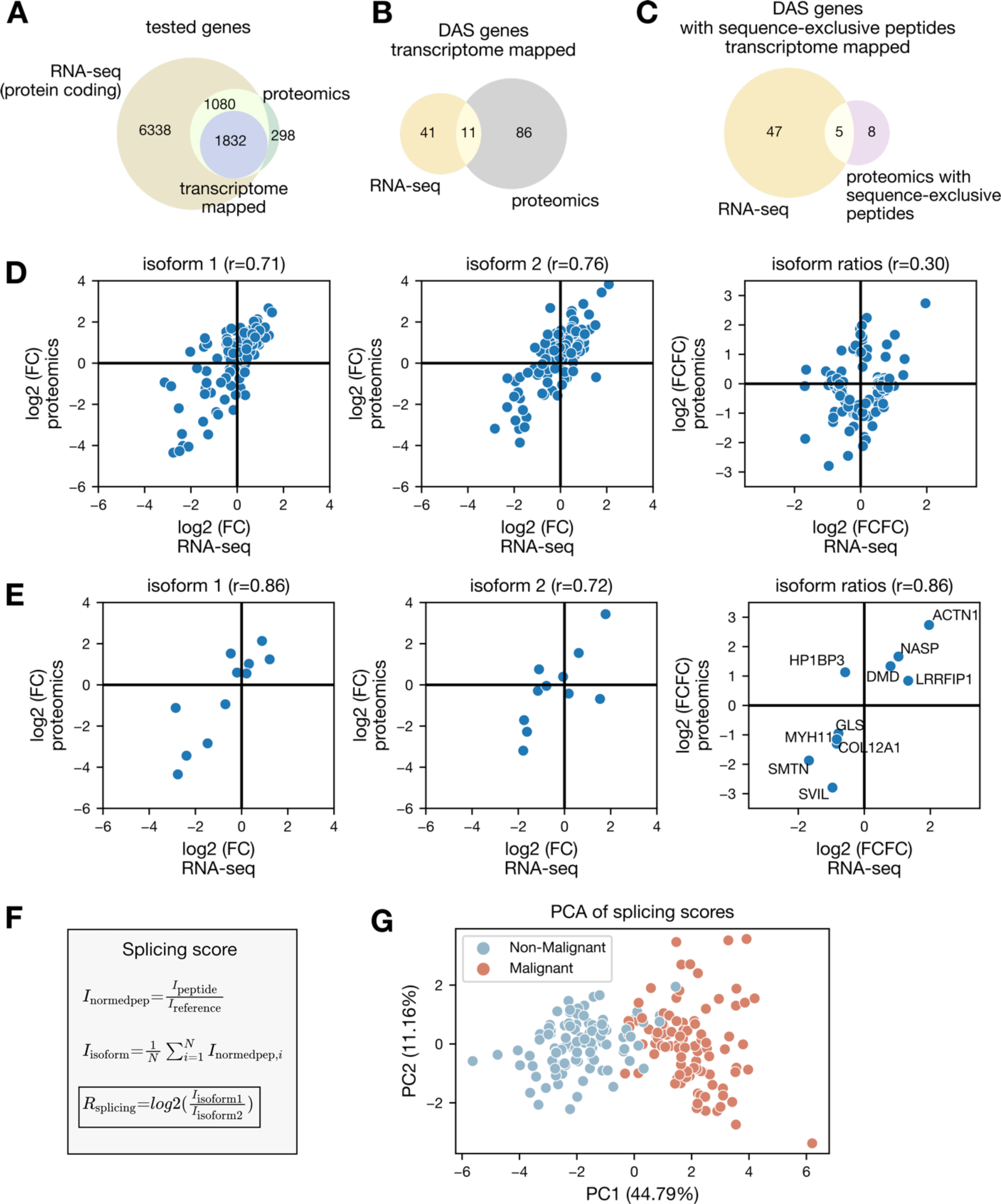
Comparing transcriptome and proteome splicing in colon cancer. A) Number of genes before and after proteome-to-transcriptome mapping in the TCGA transcriptomics and the CPTAC proteomics dataset. B) Genes classfied as DAS in both datasets. C) Genes with sequence-exclusive peptides in the proteomics data as compared to genes classified as DAS in the transcriptomics data. D) Comparison of log2 fold-changes between cancer and normal between transcriptomics and proteomics data for both isoforms (left, middle) as well as for the log2(FCFC)s between isoforms (right). All genes that are classified as DAS in either transcriptomics or proteomics data are included. E) Same comparisons as in the previous plot, subsetted to the 11 genes classified as DAS in both transcriptomics and proteomics data. F) Details of our per-gene, per sample splicing score. G) PCA of splicing scores for the samples of the entire CPTAC proteomics dataset.

#### Consistent DAS genes classify disease state

Having established that genes classified as DAS in both transcriptomics and proteomics data show high correlation for the individual isoforms as well as for the log2(FCFC)s even despite coming from different patients in different cohorts, we speculated that they might be well conserved and specific for colon cancer. As an initial check for this, we created a *splicing score* for the proteomics data which captures the degree of distinct regulation of isoforms (Figure 5F). For this score, we first needed to normalize peptide intensities, which we did here by dividing each peptide intensity through the mean intensity of the ‘ColonRef’ reference samples measured in the cohort (Methods). For each isoform, we calculated the average normalized intensity of the peptides mapping to this isoform and then determine their proportions (log2 ratios). Indeed, we did see a clear separation of normal and cancer samples when performing a principal component analysis (PCA) on the splicing score (Figure 5G). This demonstrates that not only protein intensities but also splice isoform ratios enable a characterization of disease state.

### Comparing splicing profiles for individual mice

#### Acquisition of a dataset to resolve differences between individual brain proteomes

Above, we compared cancer against normal – a case where strong regulation is expected between conditions. To investigate a converse, physiological case where differences in splicing are expected to be more subtle, we measured cortex tissue proteomes from four individual, inbred mice using data-independent-acquisition (DIA) (see Methods, Figure 6A). Three replicate measurements of each mouse were acquired, which allows to compare quantitative splicing profiles between individual mice using MS-EmpiReS’s error distributions. We selected the brain because splicing is abundant and has important functional roles in the regulation of this organ. At the same time, little is known about individual variations on the splicing level. Our single-run proteomics measurements resulted in around 50,000 sequence unique peptides and 6,000 genes measured per mouse, with few missing values (Figure 6B). To investigate splicing, we chose one mouse to serve as a reference against which we compared the other mice (see Methods). With this setup, we could test around 1,200 genes for differential splicing events (Figure 6B), of which we classified 160 as significant. Our *p_adj_* thresholds for significance were set at 0.05 for genes without sequence-exclusive peptides (i.e. peptide sequences indicating a splice event) and 0.15 for genes with sequence-exclusive peptides. These thresholds were more permissive for this analysis as in the cancer example as we wanted to observe global trends over multiple samples.

**Figure 6.**
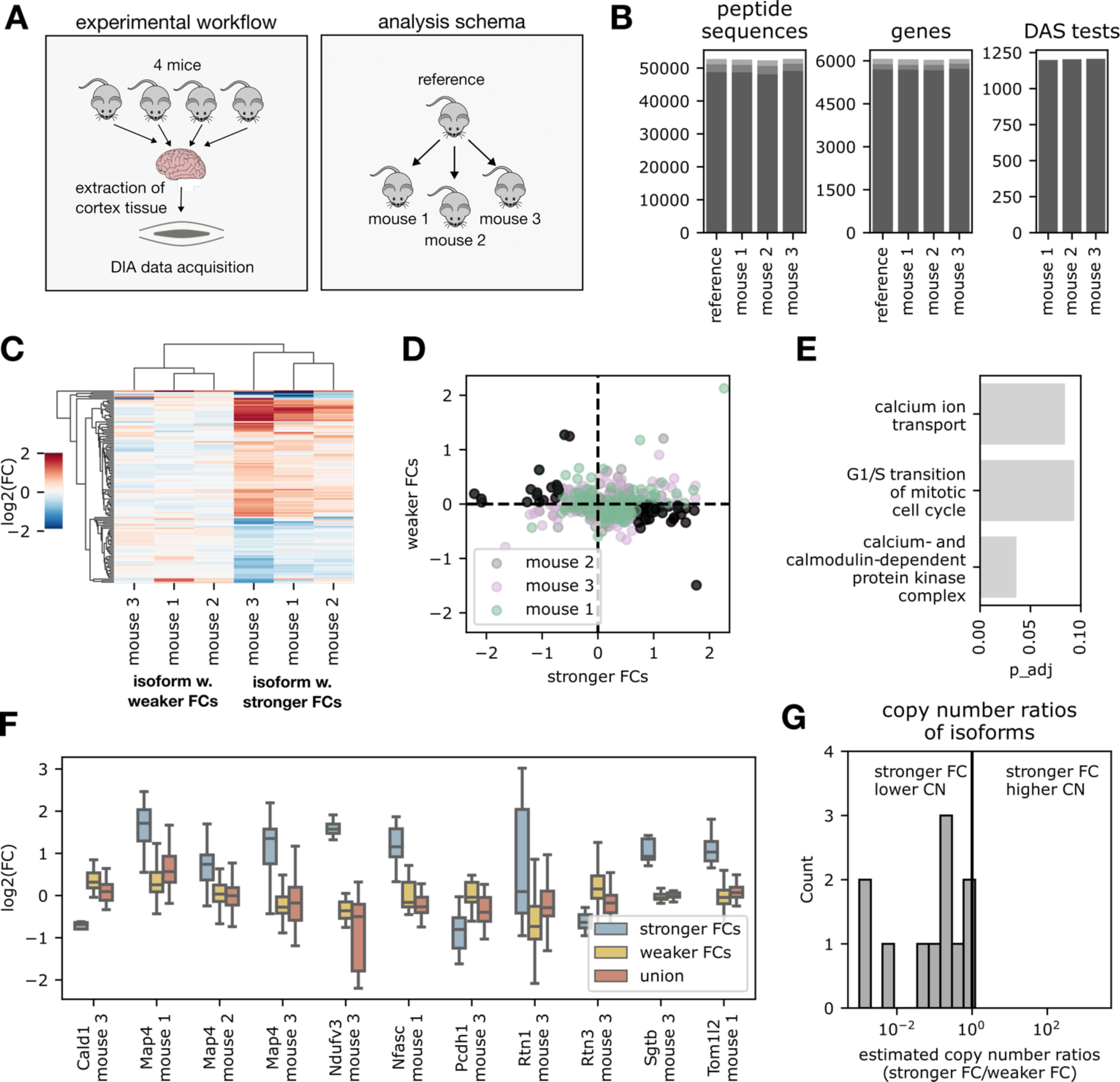
Comparing splice profiles of individual mice. A) Acquisition and evaluation scheme of DIA experiments. B) Numbers of peptide sequences, genes and DAS tests. Grey shadings indicate number of missing values (2, 1 or 0) from light to dark. C) Heat map of fold-changes relative to reference, showing two isoforms for each gene, classified as ‘strong FC’ and ‘weak FC’ isoforms. D) Scatter of the fold-changes from the stronger compared to the weaker isoforms. E) GO enrichment of the genes highlighted in black in the previous plot. F) FCs of peptides mapping exclusively to the strong FC isoform, weak FC isoform or to both isoforms for all available genes in the different mice. D) Estimated ratios between the copy numbers of the stronger FC isoform as compared to the weaker FC isoform.

#### Individuals show strongly and weakly changing isoforms

We first analyzed the regulation of all significant isoforms as a heatmap which contains the two isoforms identified by MS-EmpiReS (Figure 5C). The isoform displaying the stronger fold-change across the three mice was termed the “stronger” isoform. Our data reveal a stark contrast between the two types of isoforms, with average variances of 0.15 for the stronger and 0.05 for the weaker splicing isoforms across the 160 significantly different genes. Accordingly, the stronger isoforms separate the individuals more clearly. Comparing the fold-changes between the two isoforms directly reveals the global contrast in regulation strength, indicating that changes are driven mainly by one of the isoforms changing its abundance level (Figure 6D). We performed Gene Ontology (GO) enrichment on the genes with more strongly differing isoforms (marked grey in Figure 6D) and found “Calcium Ion Transport” to be the most enriched process. The genes associated with this process included the Calcium/Calmodulin Dependent Protein Kinases types II: CAMK2a, CAMK2b, CAMK2d. CAMK2 proteins have been shown to play key roles in learning, memory and synaptic plasticity, with functional modularity via their alternative splicing patterns which differ in a regional specific way^42–44^. Our data indicate that even individual, inbred mice can have significant differences in this therapeutically important process.

#### Variation is driven by the lower abundance isoform

To further investigate the pronounced differences between the stronger and the weaker changing isoforms we selected all genes for which we had all three sets of peptides, with one set mapping exclusively to isoform 1, one set mapping exclusively to isoform 2 and one set mapping to the union of both isoforms. Interestingly, the weaker fold-changes are very similar to the fold-changes of the union peptides with the stronger fold-changes diverging from both sets (Figure 6F). As detailed in Supplemental Text 1, this data allows us to calculate the stoichiometry of the two isoforms on the basis of the fold-changes of each set. Interestingly, the stronger isoforms are of lower abundance, indicating that the changes we see are driven by strongly responding lower abundance isoforms (Figure 6G). As the low abundance isoform still needs to be detectable by MS, real copy number differences may be larger yet in the animal.

## Discussion and Conclusion

In this study we have introduced a bioinformatics framework that very effectively detects differential alternative splicing in MS-based proteomics data based on quantitative peptide level information. MS-EmpiReS detects splicing induced differences in the relative abundance of two isoforms between conditions via fold changes of fold changes (FCFCs). Current approaches of protein-level DAS are based on directly detecting sequence-exclusive splicing in MS proteomics measurements. By their nature, these detected events are rare, which leads to a drastic loss in sensitivity upon subsequent quantification. In contrast, our approach adds additional evidence by considering many more of the measured peptides. To maximize flexibility, we specifically designed MS-EmpiReS to be a proteomics and not a proteogenomics tool that would depend on additional transcriptomics data. That said, it is easily possible to compare the proteomic and transcriptomics splicing data as shown in this study. In principle, MS-EmpiReS can analyze splicing events from any quantitative proteomics experiment where two or more conditions are compared and at least two replicates are measured in each condition. The only inputs needed are quantified peptides and a condition mapping. MS-EmpiReS comprehensively incorporates prior knowledge on splicing by screening the complete Ensembl annotation. So far, we have analyzed human and mouse data, other organisms could easily be added. With MS-EmpiReS, we provide a software package with fast execution time and simple input commands, which allows researchers to quickly screen and visualize datasets for DAS, thus a useful and easy to use addition to standard computational proteomics pipelines.

A potential limitation of MS-EmpiReS is that it is based on quantitative information underlying detected DAS events that might itself be subject to potential biases. In particular, systematic offsets in the quantification of peptides might lead to incorrect results. For example, it is possible that quantitative variations for example due to incorrect identifications, interferences or other biological effects could lead to such errors. However, on the *E. coli* benchmarking dataset we have shown well controlled false positives in a realistic biological scenario which could be hampered by all of the before mentioned biases. Our choice of *E. coli* as a test for false positive was also based on the fact that currently no benchmarking sets for DAS exist, as it is difficult to know the ground truth on splicing regulation. We suggest that the community invest in such resources. In any case, the (near) absence of false positives in our benchmarking datasets allows us to have confidence in hits in subsequent biological analyses. For additional confidence, we encourage manual inspection of the splice events via the visualizations provided by MS-EmpiReS.

We applied our method to a substantial clinical dataset, which is of intrinsic interest in cancer research, but also carries a high level of noise and also of biological variation. We were able to recover clear proteomic splice events of important regulator genes, providing novel information about the regulation of proteomic splicing in cancer. We validated our hits by comparing them to the sequence-exclusive approach, independent databases, literature knowledge as well as RNA-seq data. In a completely different scenario, we demonstrated that subtle differences in splicing can be resolved by comparing measurements of brains from individual mice to each other. This revealed individual-specific differences in splicing isoform regulation, which was driven by the less abundant isoform.

The analysis of DAS on the transcriptome level continues to generate novel insights both in basic biology as well as in clinical research^11,45–47^ despite being limited to measuring mRNA. As proteins represent the functional players in the cell, studying regulation of splicing on the protein level is crucial for obtaining further insights into the biological implications. Consequently, enabling the investigation of DAS on a proteome-wide scale should resolve many long-standing questions such as the influence of splicing on protein structure and proteome diversity. Apart from computational efforts such as MS-EmpiReS, recent developments in instrumentation promise proteomics sequence-coverage comparable to that of RNA-seq data in the near future^48^. In turn, this should elevate DAS analyses and the analyses of other types of proteoforms to a new, much more comprehensive level.

## Methods

### Simulating non spliced proteins

In the first step of the simulation, differential peptides, i.e. peptides quantified in two conditions were simulated. For each simulated peptide, *r*_1_ and *r*_2_ intensity values were drawn from a log normal distribution, with *r*_1_ and *r*_2_ being the numbers of replicate measurements in each condition. A log normal distribution is generally used as an approximation to describe protein/peptide abundances^49^. Two groups of peptides were tested against each other for each simulated protein, meaning that for each simulated protein, *g*_1_ + *g*_2_ peptides were drawn, with *g*_1_ and *g*_2_ the numbers of peptides in each group. 3,000 proteins were simulated for each test, with the parameters *r*_1_*,r*_2_*,g*_1_*,g*_2_ specified in Figure 2A.

### Benchmarking on the technical datasets

The technical dataset was downloaded as specified in the data availability section. The data was acquired in the context of a study on differential quantification^50^, where human cell lysate with differing amounts of yeast spike in were measured with LFQ and TMT-MS3. This dataset had been used in the MS-EmpiRe study^25^, where it had been processed as follows: The data was searched and quantified via MaxQuant^51^ v. 1.6.0.16 with standard settings and additional LFQ or TMT quantification set. A combined yeast (7,904 entries) and human (20,317 entries) database downloaded from Uniprot^52^ (April, 2018, reviewed) was used. For the DAS benchmark, the human proteins were selected and for each quantification method, 6 replicate runs were compared as 3 vs 3 replicates.

### Benchmarking on the *E. coli* datasets

The *E. coli* datasets were downloaded from their respective PRIDE repositories (see data availability section). For the LFQ dataset^53^, the .raw files were downloaded and searched with MaxQuant v. 1.5.7.4 against the reviewed Uniprot *E. coli* K-12 database (03/2019), using standard settings with additional “Label Free Quantification” and “Match between runs” set. For the TMT dataset (acquired via an SPS-MS3 workflow), MaxQuant search files were directly downloaded. As we collected the data from different studies, the details of the preprocessings (for example the MaxQuant versions) differed slightly. Our model is however not dependent on such preprocessings and should not be affected by this. Peptide intensities were extracted from the “peptides.txt” files for both TMT and LFQ data. Only peptides with unique mapping to a protein were considered. The conditions were compared in a pairwise manner resulting in 352 condition pairs for the LFQ data (see Supplemental Table 4) and 3 condition pairs for the TMT data. For each condition pair, we iterated through all proteins. For each protein, peptides were randomly distributed in two groups of equal size. The groups were then tested against each other for DAS, resulting in one p-value for each protein in the condition pair. Multiple testing correction was carried out using the Benjamini-Hochberg^54^ procedure on the p-values for each condition pair and proteins were classified as significant with an *p_adj_*<1% and an FCFC>1.5.

### Preprocessing the CPTAC data set

MSGF+^55^ search files and MASIC^56^ selected ion chromatograms (SICs) were downloaded from the CPTAC data portal (see data availability section). Peptides were filtered to a MSGF+ q-value <0.005, a MASIC InterferenceScore >0.9 and a PeakSignalToNoiseRatio>35. In the case of multiple identifications of the same peptide, the identification with the strongest signal was chosen. Fractions of the same TMT multiplex were merged and two normalization steps were performed. In a first step, TMT channels of the same multiplex were normalized to correct for differing sample amounts in the channels, using the MS-EmpiRe within-replicate normalization. In a second step, the TMT-10 131 channels with a technical spike-in were used for peptide specific normalization between TMT multiplexes. For each peptide, fold-changes between the reference channels were obtained and one normalization factor per multiplex was estimated by estimating the minimum error with Levenberg-Marquardt optimization. The dataset was acquired with MS2 level quantification, thereby possibly giving rise to the phenomenon of ratio compression^57^ (i.e. underestimation of absolute fold-change strength) due to co-fragmenting precursor ions. However, several factors mitigated this problem: From the data acquisition side, small isolation windows (0.7 Da) were used, which reduced the probability of co-fragmentation. Additionally, the data was highly fractionated (96 fractions of which 12 were chosen per multiplex) and the ≈200 patient samples were distributed on a total of 22 multiplexes. This should reduce the chance that the same peptide measured in multiple multiplexes has the same type of co-isolation. Rather, co-isolation would result in additional noise between replicates, which can be accurately modeled. On the computational side, we used conservative interference score cutoffs and our model is able to handle inconsistent fold-changes, as shown in the *E.coli* benchmark. As displayed in Supplemental Figure 4, our normalization results in low noise levels and separation between normal and cancer samples already on the peptide intensity level, indicating good quantification.

### Data normalization

Data were normalized similar to MS-EmpiRe^25^, which uses the concept of centralization^58^. Briefly, samples were shifted by a constant factor to cancel out systematic biases, e.g. due to differing sample amounts. The factors were derived from peptide fold-change distributions using the median for replicate normalization and the mode for normalization between conditions.

### Mapping peptides to isoforms

Peptides were mapped to genes and protein isoforms using the Ensembl homo sapiens GrCh37.75 genome annotation. Peptides mapping to more than one gene were eliminated from the analysis. In order to detect sequence-exclusive splicing, conflicting peptide pairs were searched, meaning pairs of peptides that cannot exist on the same isoform. One peptide was required to span an exon junction and the other peptide was required to be inconsistent with this junction, meaning that it starts or ends inside the junction. For the quantitative approach of MS-EmpiReS, an *equivalence class* was obtained for each peptide. Similar to the BANDITS^59^ package for transcriptomic DAS detection, we define an equivalence class as the set of all isoforms a peptide maps on. This means that peptides mapping to the same equivalence class should have equal abundances and be independent of DAS (under the assumption that the annotation is comprehensive). Peptides mapping to different equivalence classes could potentially stem from different isoforms or different mixtures of isoforms and can show abundance changes after DAS. Grouping peptides by equivalence classes hence enables very “clean” testing for DAS. In the case that conflicting sequence-exclusive peptides were detected for a gene, we grouped the remaining peptides differently and distributed all remaining peptides of the gene around the sequence-exclusive peptides. For this, we iterated through all other peptides of the gene and tested for each peptide if its equivalence class overlaps (i.e. has shared isoforms) with one sequence-exclusive peptide and does not overlap with the other sequence-exclusive peptide. If this criterion was fulfilled, we grouped it to the respective sequence-exclusive peptide. An equivalence class was included in the further analysis if there were at least two distinct peptide sequences quantified, or if a conflicting sequence-exclusive peptide was quantified in another equivalence class. To reduce the influence of technical noise in the latter case, we also required at least two peptide ions to be quantified (not necessarily with differing sequences). In total, of ≈138,000 quantified peptides, ≈64,000 mapped to equivalence classes, ≈47,000 of which passed the filtering criteria. For gene-level testing, as displayed in Figure 2, the most significant pair of equivalence classes was chosen for each gene and multiple testing correction was carried out subsequently via the Benjamini-Hochberg^54^ procedure, the corrected score is denoted as *p_adj_*.

### Calculation of FCFCs

After isoform mapping, the equivalence classes were tested against each other in a pairwise fashion. For this, all peptide pairs between two equivalence classes were obtained. For a peptide pair *p*_1_ and *p*_2_ the log2(FCFC)s were calculated. To define the log2(FCFC), we first define the peptide intensity *I*(*c,r, p*), which is dependent on condition c, replicate r and peptide p. A peptide fold-change is subsequently defined as

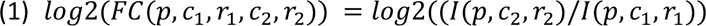

with conditions *c*_1_, *c*_2_ and respective replicates *r*_1_, *r*_2_. The log2(FCFC) is then defined as

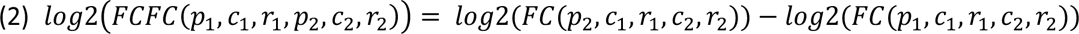

with peptides *p*_1_ and *p*_2_. We see that the only factor changing between first and second term is the peptide. In this definition, peptides *p*_1_ and *p*_2_ are only compared between identical replicates.

### Generation of empirical FCFC error distributions

To put FCFCs into a statistical context, we generated empirical FCFC error distributions from replicate measurements. As a first step, we generated empirical FC error distributions, analogous to MS-EmpiRe^25^. For this, log2(FC)s between peptides of equal sequence and charge were obtained between replicate measurements, generating an empirical error distribution of log2(FC)s. As replicate measurements carry the sum of biological and technical variation, this empirical FC error distribution should exactly reflect this variation and hence be a good estimate of the noise underlying the experiment. Analogous to MS-EmpiRe, we also separated the empirical FC error distribution into sub distributions depending on the intensities of the peptides measured.

As a general trend, peptides with lower intensity are subject to higher variation than peptides with higher intensity and it thus makes sense to have multiple empirical FC error distributions, covering different intensity ranges. To increase runtime and memory efficiency, we binned the empirical FC error distribution into log2(FC) intervals of 0.01. To generate the empirical FCFC error distributions, we took two empirical FC error distributions and created the difference distribution, as also described by Csaba et al.^41^. Technically this is simply achieved by comparing all possible pairs of bins. For each pair of bins, we calculated the log2(FCFC) by subtracting the log2(FC)s of the pair and obtained the corresponding frequency by multiplying the frequencies of the pair. Empirical FCFC error distributions were generated for all necessary pairs of intensity ranges.

### Combination of FCFCs

A pair of equivalence classes with n and m peptides generates *n*·*m* peptide pairs and each peptide pair generates a maximum of *r*_1_ ·*r*_2_ FCFCs, with *r*_1_ and *r*_2_ being the respective number of replicates in each condition. Our aim is to obtain the overall null probability for DAS of the equivalence class pair from this (possibly very large) set of FCFCs. Analogous to the MS-EmpiRe paper^25^, we transform the FCFCs into normally distributed random variables using a modified Stouffer approach and combine these variables by summation. We see that many of the *FCFC*(*p*_1_*,c*_1_*,r*_1_*, p*_2_*,c*_2_*,r*_2_) are dependent on each other. For example, the pair *FCFC*(*p*_1_*,c*_1_,1*, p*_2_*,c*_2_,2) and *FCFC*(*p*_1_*,c*_1_,1*, p*_2_*,c*_2_,3) has a shared replicate. When we consider the FCFCs to be random variables, which we combine, these dependencies affect the variance of the combined distribution and have to be taken into account. A main goal is hence to appropriately estimate the variance of the combined random variables, which can be achieved via summation over the full covariance matrix. The basic concepts for this estimation have been introduced in the MS-EmpiRe paper^25^ and a generalized version which has been used in this work, is described in detail by Csaba et al.^41^.

### Gene Ontology enrichment

Gene Ontology enrichment was based on the Gene Ontology .obo database, using the GOATOOLS python package^60^. Enrichment was calculated via overrepresentation analysis.

### Processing RNA-seq dataset

We performed transcriptomic analysis to investigate differential alternative splicing in a subset of the Colon Adenocarcinoma (COAD) cohort provided via The Cancer Genome Atlas (TCGA)^38–40^. This subset consisted of tumor and normal samples from 20 patients. Patient identifiers are listed in Supplemental Table 5. We used the mapped Binary Alignment/Matching (BAM) files provided by TCGA, which we aligned to the Ensembl homo sapiens GrCh37.75 genome annotation. The differential alternative splicing analysis was carried out using the EmpiReS^41^ software package, comparing normal against cancer tissue. The EmpiReS output tables contained the best scoring isoform pair for each gene and reported the fold-changes of the individual isoforms, the FCFC of the isoform pair and the estimated *p_adj_* for DAS. After processing, we filtered out all isoform pairs that contained non protein coding genes.

### Mapping proteomics to transcriptomics data

To align the proteomics and the transcriptomics data, we used the RNA-seq data processed with MS-EmpiReS as the reference to which we aligned the proteomics data. The reasoning for this was that the sequence coverage is in general higher for RNA-seq data, which allows higher confidence in the isoform mapping. For each isoform pair in the RNA-seq data, we then searched the pair of equivalence classes that contained both isoforms among all possible equivalence classes in the respective gene. We only considered genes as “transcriptome mapped” for which a matching pair of equivalence classes could be found.

### Comparing to reference mouse

The multi-condition mouse dataset was processed by chosing mouse with the label E1 as a reference. The other mice were compared against this reference. For each gene, the most significant isoform pair in any of the comparisons was chosen for all comparisons, allowing comparability between conditions.

### Tissue preparation

Four 4-month old C57BL/6 mice (2 male (identifiers E1, E2), 2 female (identifiers E3, E4)) were euthanatized with cervical dislocation and placed on cold surface where brain tissue was gently removed and cortex brain area was separated. The duration of the separation of the brain area was less than two minutes. Tissues were stored at -80C until further steps of preparation. All experiments were conducted following the German animal welfare legislation and regional guidelines for animal research.

### Sample preparation

Brain cortex samples (20mg) were ground to a frozen powder using a mortar and pestle in liquid nitrogen. Powdered samples were then resuspended to sodium deoxycholate (SDC) reduction and alkylation buffer (PreOmics GmbH, Martinsried, Germany) and boiled for 10 min in the thermomixer while vortexing to denature proteins. A water bath sonicator was used as next step to sonicate the lysates at full power for 30 cycles with 30 s intervals. Overnight digestion with LysC and trypsin in a 1:50 ratio (*µ*g of enzyme to *µ*g of protein) at 37°C and 1,700 rpm in the thermomixer. Peptides were acidified to a final concentration of 0.1% trifluoroacetic acid (TFA) and loaded onto StageTip plugs of styrene divinylbenzene reversed-phase sulfonate (SDB-RPS). Peptide concentrations were measured optically at 280nm (Nanodrop 2000, Thermo Scientific) and were subjected to LC-MS/MS analysis.

### Mass spectrometry analysis

LC–MS/MS analysis of tryptic peptides was performed on a quadrupole Orbitrap mass spectrometer (Orbitrap Exploris 480, Thermo Fisher Scientific, Bremen,Germany) coupled to an Easy nano LC 1200 ultra-high-pressure system (Thermo Fisher Scientific) via a nano-electrospray ion source. For each sample 200ng of peptides were separated at 600C onto a 50 cm high performance liquid chromatography (HPLC)-column (75*µ*m inner diameter,New Objective, Woburn, MA, USA; in-house packed using ReproSil-Pur C18-AQ 1.9-*µ*m silica beads; DrMaisch GmbH, Ammerbuch, Germany). The total gradient length was 180 min with increasing buffer B (80% acetonitrile and 0.1% formic acid). Full MS scans were acquired in the range of m/z 300–1,650 at a resolution of 120,000 and the automatic gain control (AGC) set to 3e6. The mass spectrometer was operated in data dependent (DDA) and data independent acquisition (DIA) mode.

### Mass spectrometry data processing

Data were processed in Spectronaut Pulsar X software (Biognosys AG, Schlieren, Switzerland, version 12.0.20491.1) using standard settings. For library construction, samples were measured by data-dependent acquisition (DDA) and computationally processed using the Pulsar Search engine. Individual samples are then measured by our DIA workflow, including matching against the library for the identification.

### Data availability

The *E. coli* proteomics data and the technical benchmarking data was downloaded from: http://proteomecentral.proteomexchange.org via the corresponding PRIDE^61^ partner repositories (LFQ E. coli: PXD000498, TMT E. coli: PXD008339, technical benchmarking: PXD007683). The colon cancer proteomics data used in this publication were generated by the Clinical Proteomic Tumour Analysis Consortium (NCI/NIH)^62^. The “PSM” data and relevant mappings were downloaded via:

https://cptac-data-portal.georgetown.edu/cptac/s/S045

The CPTAC input file for MS-EmpiReS used in this analysis is provided together with Jupyter Notebooks for figure creation as Supplemental File 2. The MS-EmpiReS outputs are provided as Supplemental File 1. The colon cancer RNA-seq data used in this publication were downloaded as bam files from The Cancer Genome Atlas (TCGA):

https://portal.gdc.cancer.gov/projects/TCGA-COAD

The sample IDs are provided in Supplemental Table 6. The EmpiReS output table is provided as Supplemental Table 8.

MS raw files of the mouse proteomics measurements as well as output information from Spectronaut have been deposited with the ProteomeXchange Consortium via the via the corresponding PRIDE^61^ partner repository and will be made available to reviewers.

## Author contributions

C.A .and G.C. designed the method with contributions from M.G. C.V. performed the wet laboratory experiments and mass spectrometry measurements. C.A, implemented the method and performed data analyses with contributions from R.Z., G.C and all authors. G.C. provided helper classes. M.M. supervised experiments, mass spectrometry measurements and data analyses. R.Z. supervised method development and data analyses. C.A., R.Z. and M.M. wrote the manuscript with contributions from all authors.

## Acknowledgements

We thank our colleagues at the chair of Bioinformatics at LMU Munich as well as in the department of Proteomics and Signal Transduction at the Max Planck Institute of Biochemistry for help and fruitful discussions, especially, Evi Berchtold, Marta Murgia, Florian Meier, Edwin Rodriguez, Ozge Karayel and André C. Michaelis. C.A. acknowledges funding from the Deutsche Forschungsgemeinschaft (DFG, Graduate School QBM). This study was supported by The Max-Planck Society for the Advancement of Science. C.A. acknowledges funding from the Bavarian State Ministry of Health and Care as a part of “DigiMed Bayern” (grant No: DMB-1805-0001). R.Z. and G.C. acknowledge support of DFG SFB 1123 Atherosclerosis.

