## Supplemental Figures for "Detecting Differential Alternative Splicing in Mass Spectrometry-based Proteomics Data"

### Supplemental Figure 1

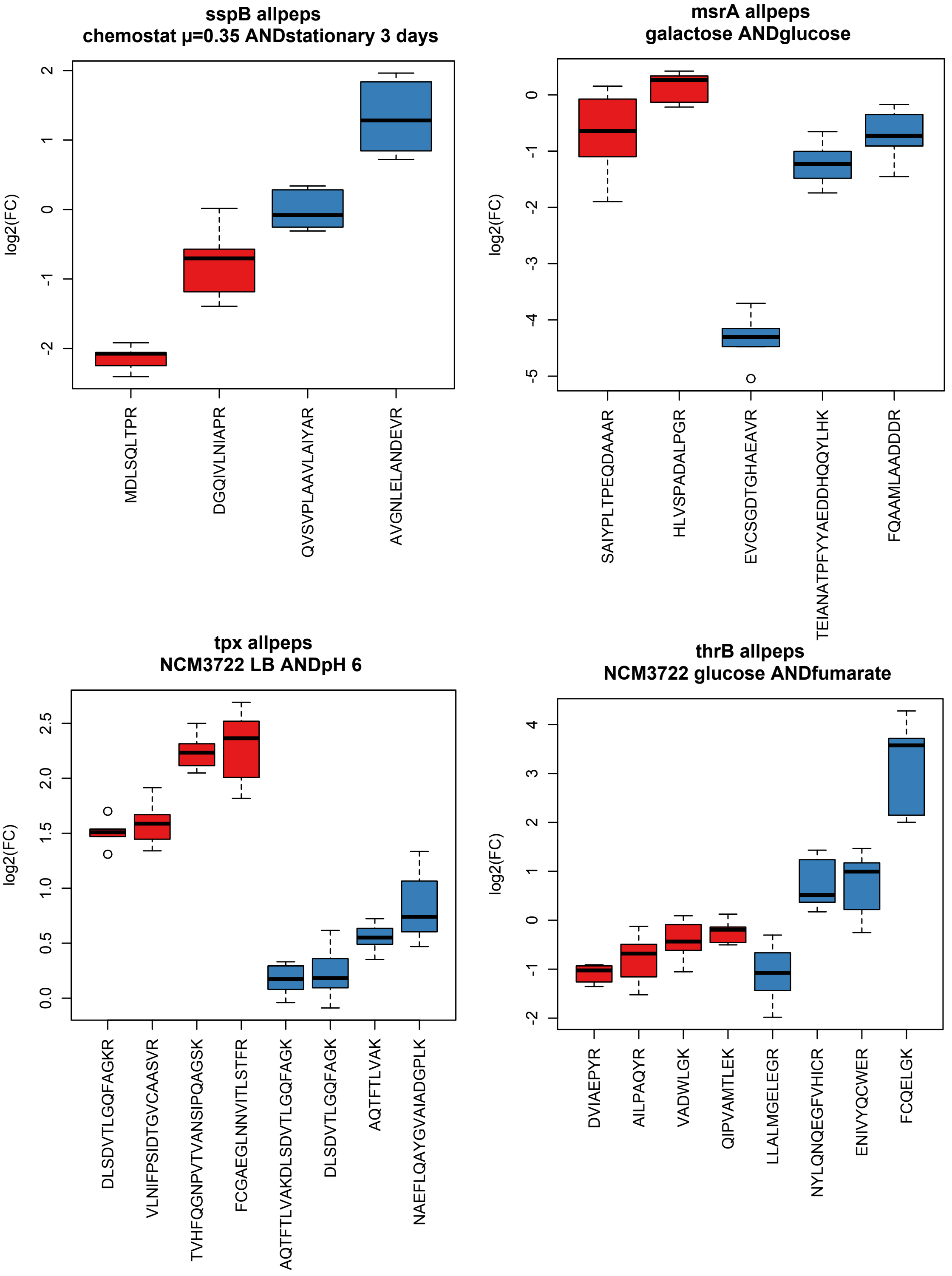


**Supplemental Figure 1** – Examples of *E. coli* proteins with inconsistent peptides. We see strong and differences in the fold changes of the individual peptides. These systematic shifts could be due to post-translational modifications or systematic biases in the data. Peptides were randomly assigned into the red and blue groups and tested with MS-EmpiReS.

### Supplemental Figure 2

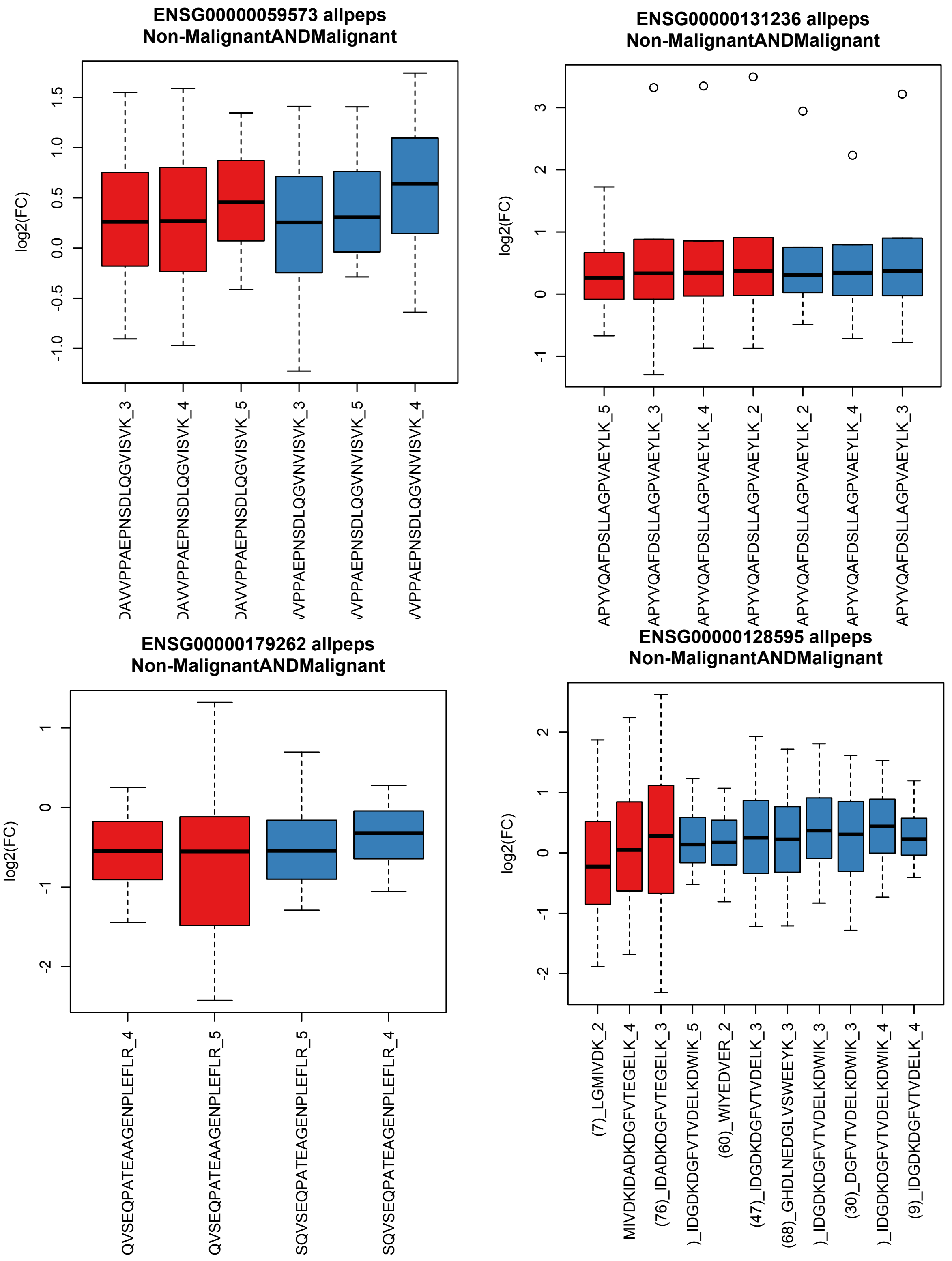


**Supplemental Figure 2** – Examples of spliced genes with no visible regulation. The genes have splice conflicts on the sequence level, however the ratios of both isoforms (red and blue) show no indication of change relative to each other between conditions.

### Supplemental Figure 3

**A**

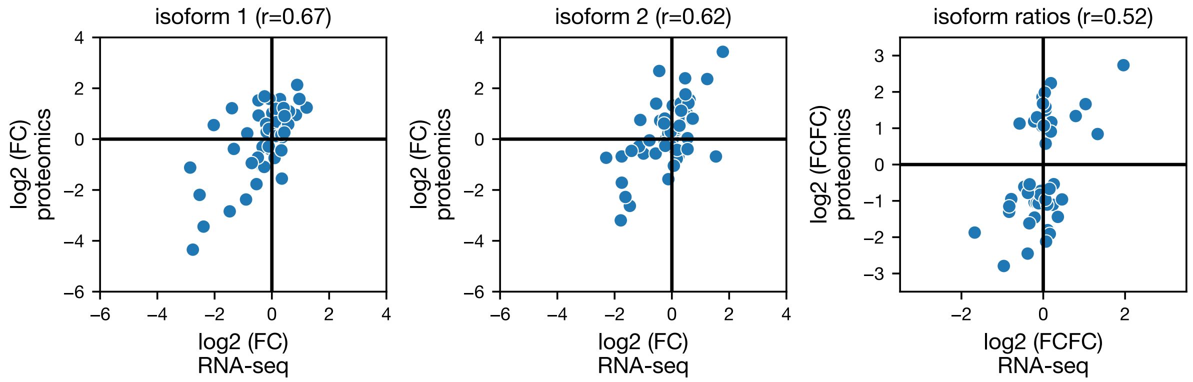


**B**

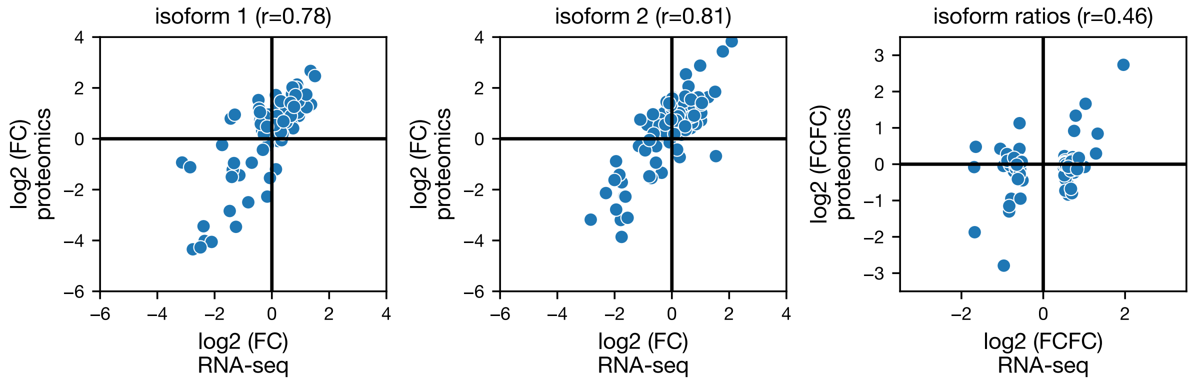


**Supplemental Figure 3** – Transcriptome to proteome correlations, analogous to Figure 5D and 5E in the main text. A) all genes that are significant in the RNA-seq data and B) all genes that are significant in the proteome data. Correlations are improved as compared to using the union of all significant genes (Figure 5D in the main text).

**Supplemental Figure 4**

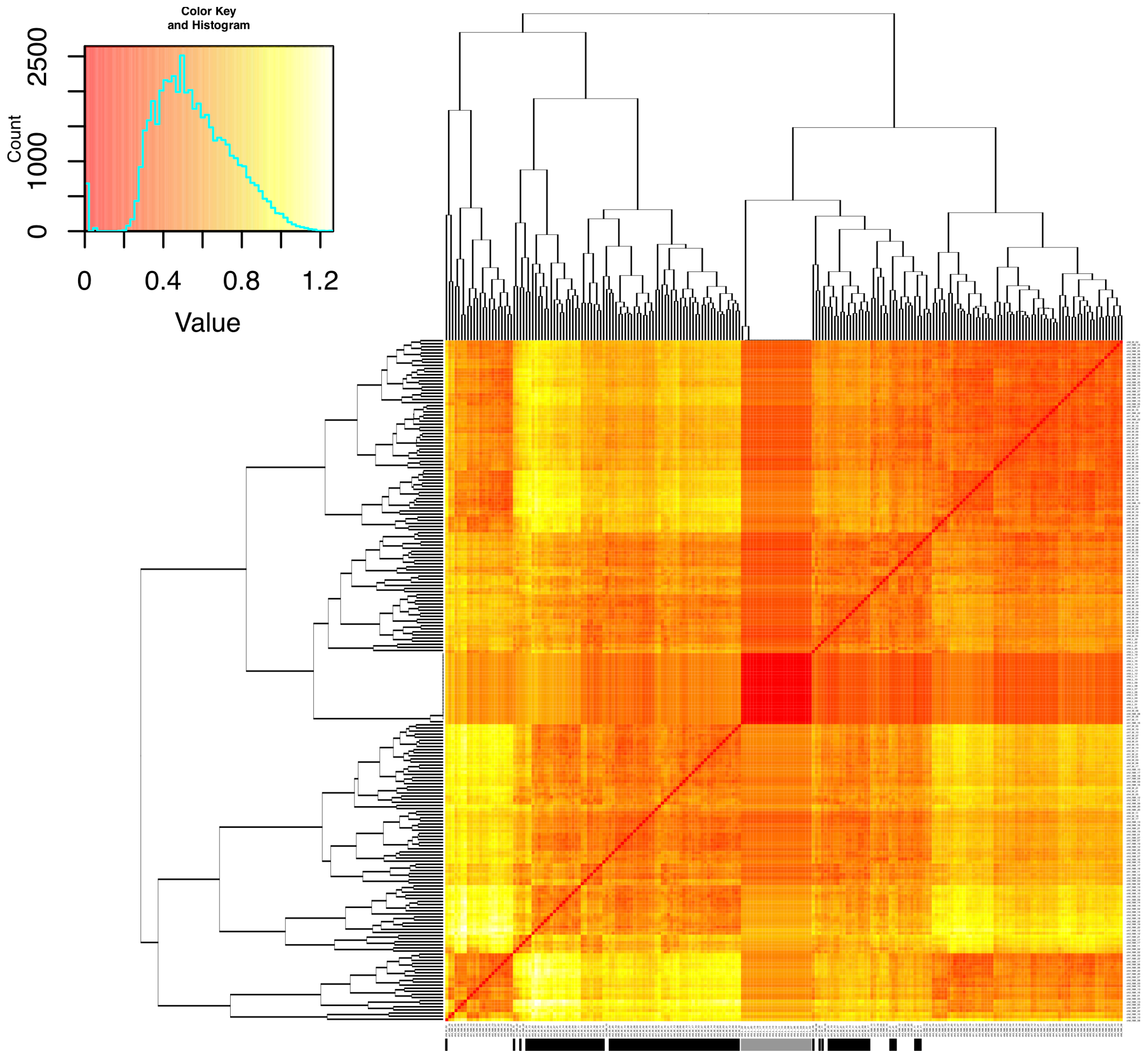


**Supplemental Figure 4** – Pairwise comparison of all samples in the CPTAC dataset after normalization. The color in the heat map indicates the standard deviation of the peptide fold change distribution between the samples. Many of the cancer and healthy samples cluster together already. Cancer samples are indicated with a black bar, 'ColonRef' channel measurements are indicated in grey.
